# Differential contribution of nitrifying prokaryotes to groundwater nitrification

**DOI:** 10.1101/2023.03.10.532121

**Authors:** Markus Krüger, Narendrakumar Chaudhari, Bo Thamdrup, Will Overholt, Laura Bristow, Martin Taubert, Kirsten Küsel, Nico Jehmlich, Martin von Bergen, Martina Herrmann

## Abstract

The ecophysiology of complete ammonia-oxidizing bacteria (CMX) of the genus *Nitrospira* and their widespread occurrence in groundwater suggests that CMX bacteria have a competitive advantage over ammonia-oxidizing bacteria (AOB) and archaea (AOA) in these environments. However, the specific contribution of their activity to nitrification processes has remained unclear. We aimed to disentangle the contribution of CMX, AOA and AOB to nitrification and to identify the environmental drivers of their niche differentiation at different levels of ammonium and oxygen in oligotrophic carbonate rock aquifers. CMX *amoA* genes accounted on average for 16 to 75% of the total groundwater *amoA* genes detected. Nitrification rates were positively correlated to CMX clade A associated phylotypes and AOB affiliated with *Nitrosomonas ureae*. Short-term incubations amended with the nitrification inhibitors allylthiourea and chlorate suggested that AOB contributed a large fraction to overall ammonia oxidation, while metaproteomics analysis confirmed an active role of CMX in both ammonia and nitrite oxidation. Ecophysiological niche differentiation of CMX clades A and B, AOB and AOA was linked to their requirements for ammonium, oxygen tolerance, and metabolic versatility. Our results demonstrate that despite numerical predominance of CMX, the first step of nitrification in oligotrophic groundwater appears to be primarily governed by AOB. Higher growth yields at lower ammonia turnover rates and energy derived from nitrite oxidation most likely enable CMX to maintain consistently high populations.

## Introduction

Nitrification is a globally relevant process of the nitrogen cycle mediated by a polyphyletic group of microorganisms [1]. The discovery of complete ammonia oxidizers (comammox bacteria, CMX) challenged the paradigm of a strict division of labor in the process of nitrification [2]. While ammonia oxidation is carried out by ammonia-oxidizing bacteria (AOB) and archaea (AOA) [3] and nitrite oxidation by nitrite-oxidizing bacteria (NOB) [2], CMX of the genus *Nitrospira* can perform complete oxidation of ammonia (NH3) to nitrate (NO_3_ ^-^) in one cell [4, 5]. CMX have been frequently reported from engineered environments such as drinking water or wastewater treatment facilities [6–8] and from natural environments like soils [9, 10], surface freshwater [11, 12] and groundwater [13, 14]. However, our knowledge about the role and relevance of CMX for nitrification activity in natural environments is still scarce.

In most environments, CMX will compete with canonical nitrifiers for either ammonia (NH3) or nitrite (NO ^-^) or both, raising the question of the actual contribution of CMX to nitrification processes and of the factors that control their co-occurrence with AOB, AOA, and NOB. Recent investigations consider affinity to NH3, growth rates, growth yields, and metabolic versatility as key factors that determine the niche differentiation and competition between CMX, AOB, and AOA [2, 15]. Since CMX exhibit higher affinity to NH3 and oxygen than terrestrial AOB [15, 16] along with higher growth yields compared to AOB and AOA [2], CMX might have a competitive advantage over AOB in environments that are limited in NH3 and ammonium (NH ^+^) such as oligotrophic groundwater. Particular lineages of AOB and AOA exhibit adaptation to NH_4_ ^+^ limited conditions [17–20] and have previously been reported from groundwater environments [21–23]. As such, oligotrophic groundwater represents an excellent microbial habitat to study the co-occurrence of CMX, AOB and AOA and their contribution to nitrification activity.

Based on the phylogeny of the ammonia monooxygenase (Amo), the key enzyme of ammonia oxidation, CMX *Nitrospira* are divided into two clades, A and B [2, 4]. Metagenomic studies uncovering the phylogeny and metabolic features of CMX proposed an ecophysiological niche differentiation between the two clades in natural habitats [4, 21, 24–26]. Those findings are linked to genomic variability in nitrogen uptake and alternative energy metabolism [25, 26], and several studies suggest a more prominent role of clade A in nitrification. So far, CMX enrichments and isolates comprise only clade A representatives [27]. ^13^C stable isotope probing confirmed growth of CMX clade A with NH_4_ ^+^ during incubations of wetland coast sediments [28], and CMX clade A were highly abundant in salt marsh sediments showing strong potential for nitrification [29]. However, the ecophysiological niche differentiation between the clades has rarely been studied in the context of their co-occurrence and competition with canonical nitrifiers.

Previous studies from karstic carbonate rock aquifers of the Hainich Critical Zone Exploratory in central Germany pointed to strong links between *in situ* nitrification and chemolithoautotrophic carbon fixation [30, 31]. In addition, genome resolved metagenomics revealed an important role of CMX *Nitrospira* in the nitrogen and carbon cycle in this oligotrophic habitat [21]. The aquifers are characterized by a high spatial heterogeneity of NH_4_ ^+^ and NO_3_ ^-^ concentrations across an oxygen gradient, linked to differences in groundwater flow regime and distance to groundwater recharge areas [28, 29, 30]. These settings are ideal for the investigation of the ecophysiological niche differentiation of CMX clade A and B, AOB and AOA.

We combined ^15^N-based nitrification rate measurements with selective nitrification inhibitors and genome-resolved metagenomics and metaproteomics to assess the contributions of CMX *Nitrospira*, AOB and AOA to overall groundwater nitrification in these aquifers and to follow the composition of the groundwater nitrifier populations across the hydrochemical gradients. We hypothesized that (i) CMX act as key ammonia oxidizers in these oligotrophic groundwater habitats and account for a large fraction of overall nitrification activity. We further hypothesized that (ii) distribution patterns of ammonia oxidizers across gradients of NH_4_ ^+^ and oxygen availability are linked to their ecophysiological niche differentiation, indicated by differences in encoded metabolic potentials.

## Materials and methods

### Study site characteristics and sampling of groundwater

Groundwater was obtained from karstic limestone aquifers at the Hainich Critical Zone Exploratory (CZE) in Thuringia (Germany) within the framework of the Collaborative Research Center (CRC) AquaDiva [34]. Further details about location and geological setting are reported elsewhere [33, 34]. Sampling wells H14, H32, H42, H43, H52 and H53 at varying depths provide access to oxygen-deficient (less than 15 µM DO) groundwater from an upper mudstone-dominated low-flow aquifer (HTU) and wells H13, H31, H41 and H51 (mean values 154 to 672 µM DO) provide access to oxic groundwater from a lower fast-flow limestone-dominated karstified aquifer (HTL) [32, 34] (Figure 1A, Figure 1B). The main recharge zone for groundwater is located at the hilltop of the transect near wells H13 and H14. Groundwater collected from regular monthly sampling between 2018 to 2021 was obtained using submersible motor pumps (Grundfos, Denmark). Hydrochemical parameters (pH, T, TOC, DOC) were determined as described in [32, 33]. NH_4_ ^+^ (sum of NH and NH ^+^), NO_2_ ^-^ and NO_3_ ^-^ concentrations were measured with colorimetric methods [35, 36] from groundwater filtered through 0.2 µm-pore size sterile polyvinylidine fluoride filters. . Concentrations of free NH3 were calculated based on pH and NH_4_ ^+^ concentrations (pKa NH ^+^/NH_3_ = 9.73 at 10°C). Groundwater for molecular analysis was collected in sterile 10 L fluorinated polyethylene containers and filtered through 0.2 µm polycarbonate filters (Nuclepore, Whatman). Additional groundwater samples for metaproteomic analysis were collected from wells H14, H41, H43 and H52 in January 2019 and filtered through 0.2 µm pore size filters (Merck Millipore, Germany). All filters were immediately frozen on dry ice and stored at -80 °C until extraction.

**Figure 1:**
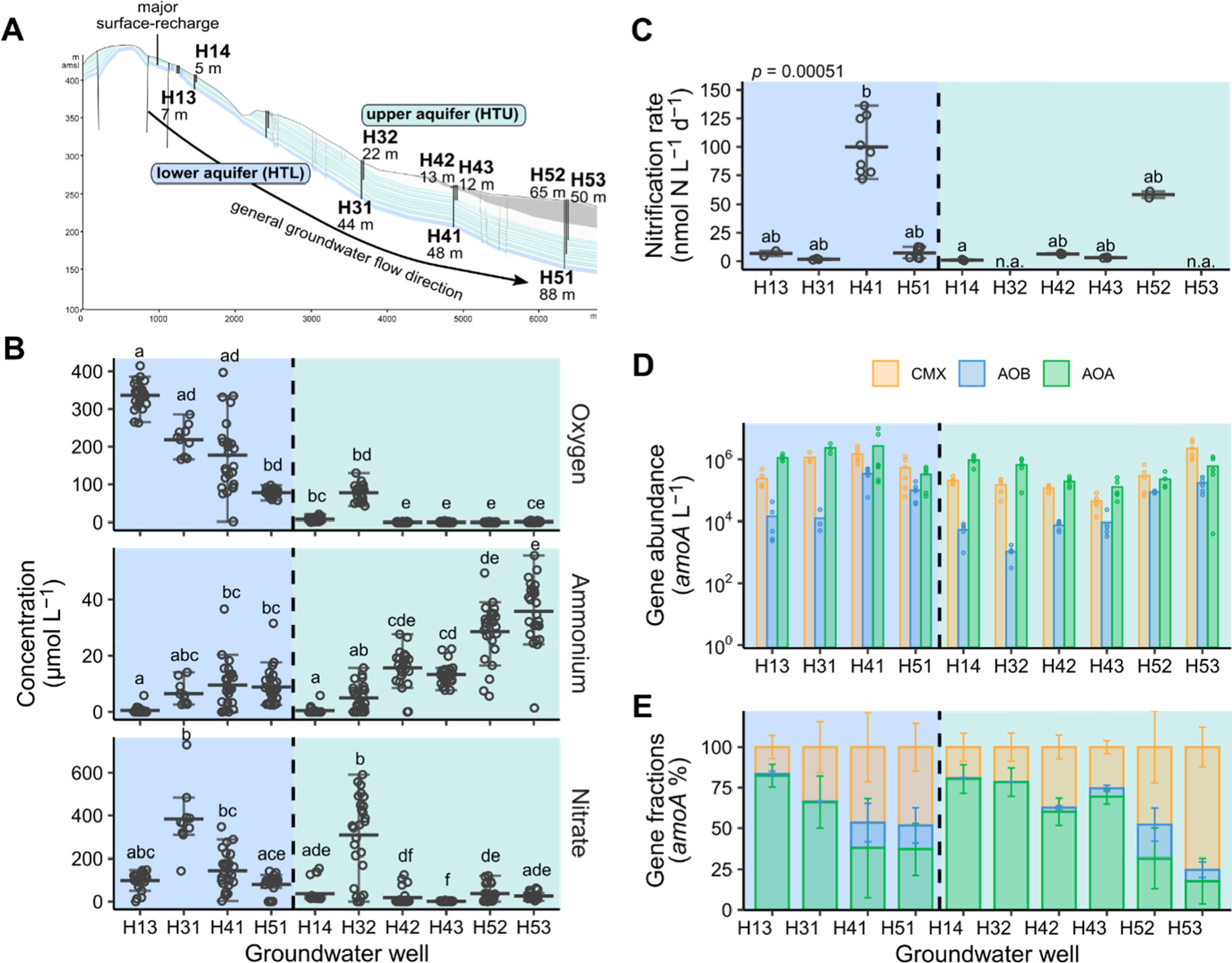
**A)** Schematic cross-section of the Hainich groundwater monitoring transect showing two superimposed aquifer assemblages accessible by 10 wells with various depths. **B)** Overview of the concentrations of dissolved oxygen, ammonium, and nitrate from oxic (blue) and hypoxic/ anoxic (green) groundwater wells (n = 24 per well). Single dots represent one measurement, the crossbar indicates the mean of all values, and the error bars show the variation from the mean. Outliers are beyond the error bars. Lowercase letter code indicates significant differences based on Dunn’s multiple comparison test (*p* < 0.05). **C)** Nitrification rates assessed by ^15^N based labeling of groundwater from five days incubation close to in situ conditions (n ≥ 3 per well; n.d. = not detected; n.a. = not available). **D)** Gene abundances of *amoA* per L and **E)** fractions of *amoA* genes in groundwater from different groups of ammonia-oxidizing prokaryotes in 10 groundwater wells (n ≥ 3).

### 15N-isotope labelling and rate measurements

Groundwater was collected in sterile 0.5 L glass bottles and filled from the bottom using sterile tubing. Bottles were overfilled with three volumes exchanges before they were capped with silicone septa without headspace to minimize alterations of *in situ* oxygen concentrations. At each sampling campaign, groundwater was collected in triplicate bottles for each well along with one control and kept at 4°C until further processing within 2 hours. Prior to the ^15^N labelling, 10 ml of each sampling bottle was removed for analysis of inorganic nitrogen concentrations and pH, and replaced with N2. Bottles containing anoxic groundwater were amended with 1 % sterile oxygen (4 ml 100% oxygen) to stimulate nitrification under oxygen-deficient conditions. Groundwater of control bottles was sterile filtered through 0.2 µm filters (Supor, Pall Corporation, USA). Sterile filtered ^15^N-labeled ammonium sulfate solution ((^15^NH4)2SO4) (98 at%, Cambridge Isotope Laboratories, Tewksbury) as the substrate source for nitrifiers was added to all 0.5 L bottles to a final concentration of 50 µM, at a natural background of 0.5 to 55 µM NH_4_ ^+^. Nitrification rate measurements were conducted according to Overholt *et al.* [21] and detailed descriptions are provided in Supplementary Methods.

### Incubation experiments in the presence of nitrification inhibitors

The ammonia and nitrite oxidation activity of CMX, and the nitrite oxidation activity of NOB were selectively inhibited by 1 mM potassium chlorate (KClO3) [7], and 100 µM allylthiourea (ATU) was used to inhibit the ammonia oxidation activity of CMX and AOB [7, 37]. Nitrification rate measurements with ^15^N tracer were carried out over five days as described above. Groundwater from well H41 was collected in February 2020 and amended with 50 µM NH_4_ ^+^ as (^15^NH4)2SO4 and one of the nitrification inhibitors along with untreated controls.

Groundwater for mesocosm experiments was collected in March and May 2021 and was filled in sterile 2 L glass bottles without headspace to assess potential growth of ammonia oxidizers with NH_4_ ^+^ in the presence of nitrification inhibitors at the same concentrations as before. Groundwater from oxic well H41 (10 µM NH_4_ ^+^ *in situ*) was adjusted to 10 and 50 µM NH_4_ ^+^, and groundwater from hypoxic well H53 (36 µM NH_4_ ^+^ *in situ*) was used without additional supplement. All samples were incubated for 15 to 25 days at 15°C without agitation in the dark. Subsamples for nitrogen compound analysis were taken at the onset and every two to three days until the end of the incubation to monitor changes of the nitrogen chemistry (Supplementary Methods). Incubation of well H41 with an initially 10 µM NH4^+^ was adjusted again to 15 µM NH_4_ ^+^ after 15 days. After incubation, the groundwater was filtered on 0.2 µm polycarbonate filters and stored at -80°C for later extraction of nucleic acids.

### Nucleic acid extraction, cDNA-synthesis and quantitative PCR

Genomic DNA and total RNA of microbial biomass from the 0.2 µm filters were extracted using the DNeasy PowerSoil kit (Qiagen, Germany) and the RNeasy PowerWater kit (Qiagen, Germany), respectively, according to the manufacturer’s specifications. RNA was further processed with the TURBO DNA-free kit (Thermo Fischer Scientific, Germany) to remove remnant DNA in the samples before cDNA synthesis [38]. Reverse transcription was performed using the GoScript Reverse Transcription System (Promega, USA). Gene and transcript abundances of *amoA* from CMX *Nitrospira*, AOB, AOA and of *nxrB* from *Nitrospira*- like NOB were determined using a CFX96 qPCR cycler (Bio-Rad, USA). Primer pairs for quantification were comamoAF/ comamoAR for CMX *amoA* [11], amoA-1F/ amoA-2R for AOB *amoA* [39], Arch-amoAF/ Arch-amoAR for AOA *amoA* [40] and nxrB169f/ nxrB638r for *Nitrospira*-like *nxrB* [41]. More details are provided in the Supplementary Methods. For testing of potential correlations between CMX *amoA* and Nitrospira *nxrB* genes, we assumed on average one copy of *amoA* and 1.5 copies of *nxrB* per cell. Calculation of the correction factor 1.5 was based on the analysis of putative canonical *Nitrospira*-affiliated MAGs from the Hainich groundwater metagenome, which harbored 1 to 2 *nxrB* gene copies per genome [21]

### Amplicon sequencing of *amoA* and 16S rRNA genes and sequence analysis

Amplicon libraries of CMX, AOB and AOA *amoA* genes were created using the NEBNext Ultra DNA Library Prep Kit for Illumina (New England Biolabs, MA) according to the manufacturer’s guideline. The same primers as for qPCR were used for CMX and AOB *amoA*, while primers Arch-amoA-104F/ Arch-amoA-616Rmod [42, 43] were chosen for sequencing of AOA *amoA* genes due to shorter fragment length (Supplementary Methods). Sequencing was conducted on an Illumina MiSeq using v3 chemistry. After trimming primer sequences, raw reads were first quality filtered with DADA2 1.22.0 [44] and further processed with Mothur v.1.46.1 [45] (Supplemental Methods). OTUs were assigned at 95% sequence identity for CMX, 95% for AOB [46] and 96% for AOA [42] (see Dataset S3).

Amplicon libraries of the bacterial 16S rRNA gene were constructed as described elsewhere [47] to assess the relative abundance of nitrite oxidizers present in the groundwater, including non-*Nitrospira* NOB. Raw sequence reads were processed using DADA2 1.22.0. Taxonomy was assessed using the SILVA reference data base release 138.1 [48]. Sequence analysis was performed using the R package phyloseq [49].

### Inferring metabolic potential from metagenome assembled genomes of CMX *Nitrospira* and other nitrifiers

To assess metabolic capabilities of the two CMX clades, genomic features of metagenome assembled genomes (MAGs) were investigated. Groundwater sampling for metagenomic sequencing and generation of MAGs were described in detail in [21, 50]. Here, Hainich groundwater MAGs, which were taxonomically classified as *Nitrospira* on genus level and 243 *Nitrospira* reference genomes downloaded from NCBI (July 2021) were aligned and screened for both *amoA* and *nxrB* genes with HMMER in Anvi’o v7.1 [21] using custom AmoA and NxrB databases containing sequences from the FunGene repository [51].

After sorting good quality MAGs (> 90 % completeness, except *N. inopinata* with 42.25 %) and dereplication, 35 CMX *Nitrospira* genomes were kept for final analysis. The affiliation of those genomes to CMX clade A and clade B was achieved by alignment of protein sequences of bacterial single copy core genes detected with default ‘Bacteria_71’ hmm profile using MUSCLE [52] in Anvi’o and subsequent construction of a maximum likelihood phylogenetic tree using FastTree with 1000 bootstrap iterations. The tree was then processed with iTOL for visualization [53]. The genomic repertoire of representative groundwater nitrifier MAGs was inferred by KEGG orthology (KO) previously generated using kofamscan [21] (Supplementary Methods). Relative transcriptional activity of each investigated MAG was extracted from Overholt *et al.* [21], who determined the transcriptomic coverages for each open reading frame from each MAG from previously generated metatranscriptomic libraries of the groundwater [30].

### Metaproteomic analysis

Proteins were extracted from filters using a phenol-chloroform based protocol as previously described [54]. Proteins were subjected to SDS polyacrylamide gel electrophoresis, subsequently in-gel tryptic cleavage and LC-MS/MS analysis was performed as previously described [55] (Supplemental Methods). Translated amino acid sequences from all genes of the MAGs obtained from the metagenomic datasets [21, 50] were used as a reference database for protein identification. Taxonomic and functional annotations of identified peptides were transferred from the metagenomic dataset [21, 50]. Mass spectrometry proteomics data have been deposited into the ProteomeXchange Consortium via the PRIDE [30] partner repository.

### Statistical analysis

All statistical analyses were conducted using R 4.12 (R Core Team, 2020) and packages FSA [57], ComplexHeatmap [58] and vegan [59]. Analysis details are explained in the Supplementary Methods. To analyze communities of CMX, AOB and AOA together, the absolute *amoA* read abundances of OTUs > 2% of all reads for each ammonia oxidizer community were determined as followed: The percentage of each OTU was calculated from the relative abundance obtained by *amoA* amplicon sequencing and was multiplied by the total *amoA* counts from corresponding samples measured with qPCR.

## Results

### Nitrification activity and distribution of nitrifiers across the different groundwater wells

Nitrification rates in groundwater varied over two orders of magnitude ranging from 0.54 – 136 nmol N L^-1^ d^-1^ across the wells (Figure 1C). We observed the highest rates at oxic well H41 with 93 ± 24 nmol N L^-1^ d^-1^ and high activity at hypoxic well H52 with 58 ± 4.0 nmol N L^-1^ d^-1^. Rates were positively correlated with NH_4_ ^+^ concentrations in the groundwater (*r*_s_ = 0.67, *p* < 0.05), which were on average below 40 µM and significantly higher in the hypoxic wells (Figure 1B). NO_3_ ^-^ concentrations ranged from 1.7 to 38 µM in hypoxic and from 80 to 384 µM in oxic groundwater and showed no correlation with the rates across the transect.

CMX *amoA* genes accounted for 16 ± 7.2 to 75 ± 12.3 % of the total groundwater *amoA* genes detected (mean values per well, see Dataset S2). Proportions of CMX *amoA* and AOB *amoA* genes increased substantially to maxima of 95% and 34% at wells with higher NH_4_ ^+^ concentrations (H41, H51, H52, H53), and gene abundances were positively correlated to NH_4_ ^+^ (*r* = 0.48, *p* < 0.01; *r*_s_ = 0.44, *p* < 0.01), respectively (Figure 1E, Figure 2D). In contrast, AOA dominated the ammonia-oxidizing community by up to 90% in wells with lower NH_4_ ^+^ concentrations located closer to the main recharge area (H13 and H14). Oxic well H41 and anoxic well H53 harbored the highest absolute *amoA* gene and transcript abundances of CMX and AOB (Figure 1D, Figure S 1A). Nitrification rates were strongly correlated with AOB *amoA* gene abundances (*rs* = 0.87, *p* < 0.001) (Figure 2D), and AOB also had the highest *amoA* transcript/ gene ratios across all wells (Figure S 1B).

**Figure 2:**
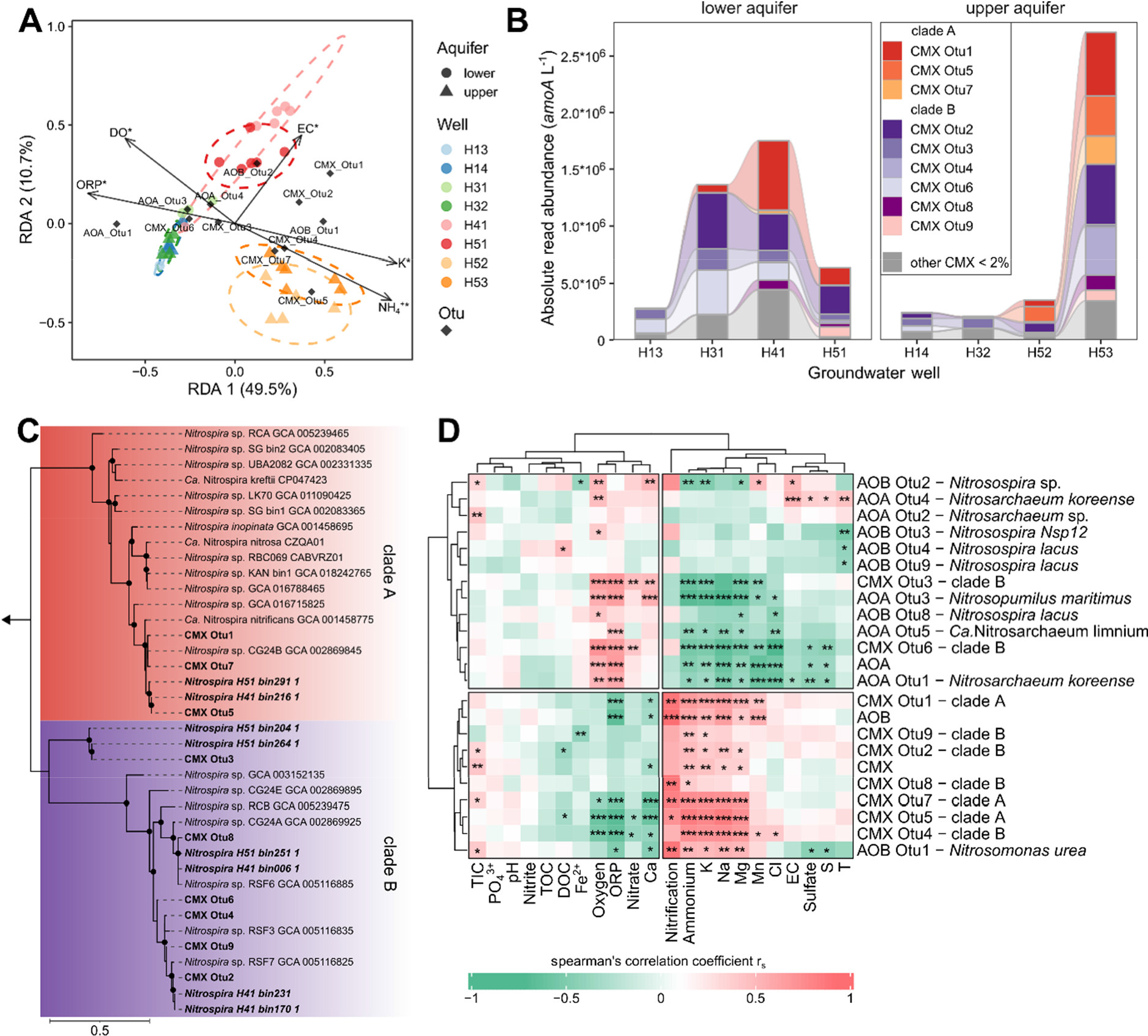
**A)** Redundancy analysis (RDA) demonstrates association between distribution of ammonia-oxidizing community (relative *amoA* read abundances of CMX, AOB and AOA OTUs which account for < 2% sequence reads of each community were normalized to *amoA* gene abundances to generate absolute *amoA* read abundances) and significant hydrochemical parameters fitting (*p* < 0.05) with nitrifier community distribution across eight groundwater wells (n ≥ 3). Dotted ellipses show the confidence of clusters assuming a multivariant normal distribution. **B)** Community composition of CMX *Nitrospira* showing dominant OTUs (< 2% of sequence reads) affiliated to clade A and clade B across eight groundwater wells (n ≥ 3 per well). Each bar displays the mean absolute *amoA* read abundance of OTUs. **C)** Maximum likelihood phylogenetic tree of deduced AmoA amino acid sequences of CMX *Nitrospira* depicting evolutionary relationship between reference genomes, and sequences of OTUs and MAGs from Hainich groundwater. The tree was constructed using the JTT substitution model with gamma distribution and 1000 bootstrap iterations. Bootstrap support values greater than 75% are denoted with black dots. An AmoA sequence from *Nitrosomonas oligotropha* served as outgroup indicated by the arrow. **D)** Pairwise correlations between total *amoA* gene abundances of CMX, AOB and AOA, absolute *amoA* read abundances of OTUs and groundwater hydrochemical parameters (TIC = total inorganic carbon, TOC = total organic carbon, DOC = dissolved organic carbon, ORP = redox potential, EC = electrical conductivity) using Spearman’s rank correlation. Hierarchical clustering is based on Euclidean distance. Color code of correlation coefficient rs displays strength of association using following significance levels: * *p* < 0.05, ** *p* < 0.01, *** *p* < 0.001.

Across all wells, abundances of CMX *amoA* genes and *Nitrospira nxrB* genes encoding nitrite oxidoreductase were positively correlated to each other (*rs* = 0.72, *p* < 0.001) (Figure S 1D). Assuming one *amoA* and 1.5 *nxrB* copies per genome (see Dataset S4), CMX accounted for 18 to 44 % and 48 to 70 % of the *Nitrospira* population at oxic well H41 and hypoxic well H52, respectively (Figure S 1C), which exhibited the highest observed nitrification rates (Figure 1C).

Furthermore, CMX and *Nitrosomonadaceae* had the highest relative abundance of peptides at wells H41 and H52 (Figure S 2). Out of the *Nitrosomonadaceae* affiliated peptides, 88.3% were assigned to *Nitrosomonas*.

The ammonia oxidizer community composition showed formation of clusters depending on local groundwater chemistry and the location in the transect (Figure 2A). The CMX community was dominated by three OTUs affiliated to clade A and six OTUs affiliated to clade B, of which clade A OTUs were closely related to *Ca.* Nitrospira nitrificans (Figure 2C). These clade A OTUs increased in abundance at wells H41, H51, H52 and H53 and accounted for up to 50% of the CMX community (Figure 2B). CMX clade A affiliated OTUs were also positively correlated to the nitrification rates (*rs* = 0.72 – 0.88, *p* < 0.05), while only two low abundant clade B affiliated OTUs correlated with the rates (Figure 2D). The AOB community was mainly dominated by AOB OTU1 affiliated to *Nitrosomonas ureae*, which was the only AOB phylotype showing a strong correlation with nitrification rates (*rs* = 0.88, *p* < 0.01). The most dominant AOA OTU1 related to *Nitrosoarchaeum koreense* accounted for an average of 70% of all AOA across the groundwater sites (Figure S 3). Clustering based on the correlation with hydrochemical parameters suggested similar ecological preferences of (i) AOA and some clade B affiliated CMX OTUs, and (ii) *Nitrosomonas ureae* affiliated AOB OTU1 and CMX clade A affiliated OTUs (Figure 2D). Among potential nitrite oxidizers, *Nitrospira* sp. were consistently present in all the samples, while 16S rRNA gene sequences affiliated with *Nitrospinaceae*, *Cand*. Nitrotoga, and *Nitrobacter* occurred only occasionally, and relative abundances of these groups were at least one order of magnitude lower than those of *Nitrospira* (Figure S 4).

### Metabolic capabilities of CMX *Nitrospira* clades and AOB

We analyzed nine CMX affiliated metagenome assembled genomes (MAGs) from the Hainich groundwater [21], which were phylogenomically affiliated to two CMX clade A MAGs and seven clade B MAGs (Figure 3A). Both clade A MAGs (H51-bin291-1 and H41-bin216-1) were closely related to *Ca.* Nitrospira nitrificans on the genome level and showed 99% amino acid sequence identity to the AmoA of CMX OTU5 (Figure 2C). AmoA sequences of Clade B representative MAGs showed closest relation to AmoA of CMX OTU2, OTU3, OTU8 and OTU9.

**Figure 3:**
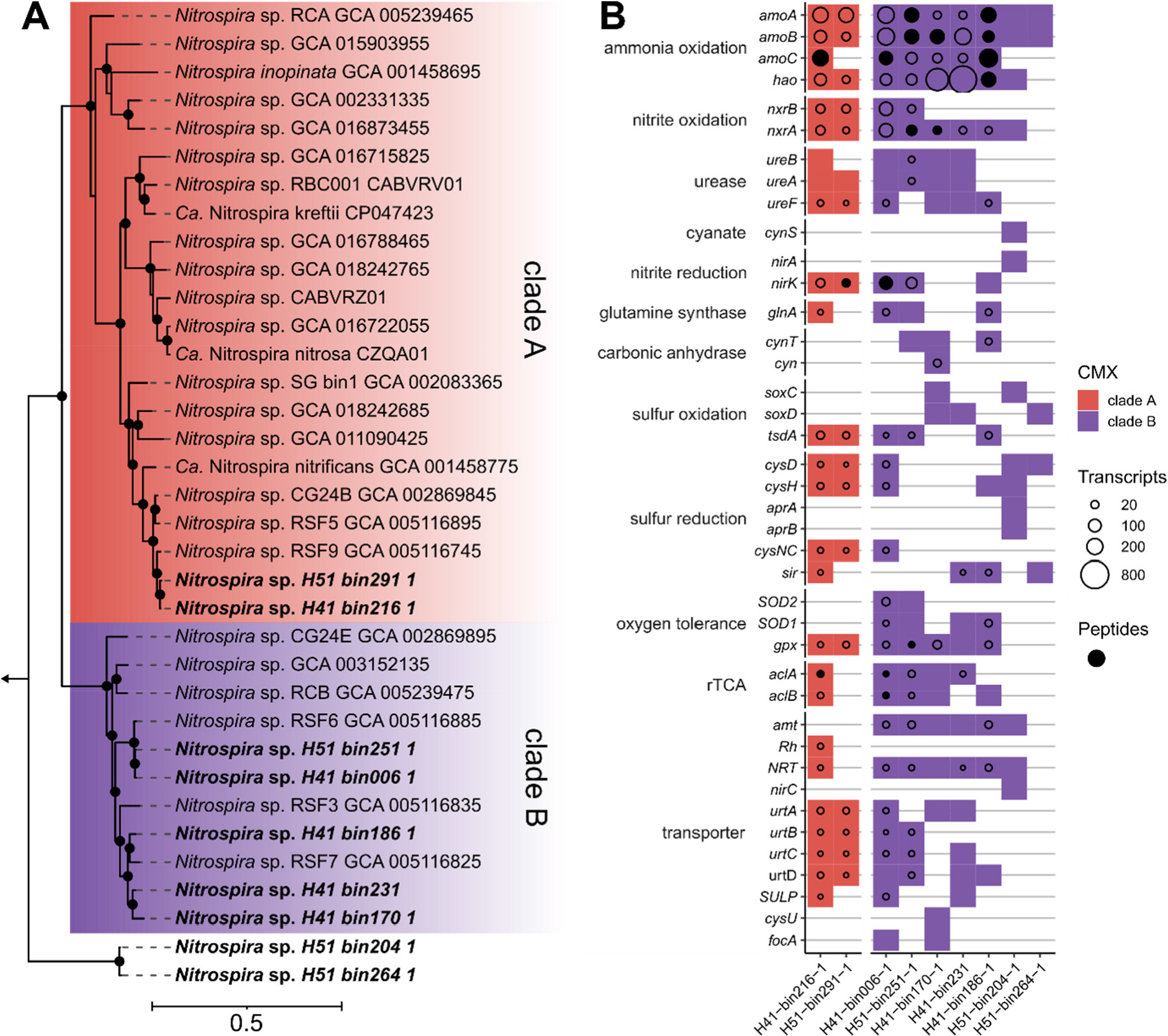
**A)** Maximum likelihood tree of bacterial core protein sequences of CMX *Nitrospira* from nine MAGs originating from Hainich groundwater depicting the evolutionary relationship to reference genomes. The tree was constructed using the JTT substitution model with gamma distribution and 1000 bootstrap iterations. Bootstrap support values greater than 75% are shown as black dots. AmoA sequences from *Nitrosomonas oligotropha* and *Nitrosomonas europaea* served as an outgroup indicated by the arrow. **B)** Metabolic capacities of nine groundwater CMX *Nitrospira* MAGs inferring genomic potential for nitrogen and sulfur energy metabolism, and corresponding transporter types as well as protective systems against radical oxygen species, and genes involved in carbon fixation (rTCA = reductive tricarboxylic acid cycle). Sizes of circles display the transcript coverage of each gene mapped from groundwater metatranscriptome. If more than one gene was present, maximum transcriptional coverage is shown. Filled circles indicate peptides from metaproteomic data matching for displayed genes. rTCA = reverse tricarboxylic acid cycle.

Genes encoding essential components for ammonia oxidation, nitrite oxidation and urea degradation were shared in MAGs representing both CMX clades (Figure 3B) and most of these genes were highly transcribed in the groundwater. Metaproteomic analysis revealed the presence of peptides related to AmoA, AmoB, AmoC, Hao and NxrA of *Nitrospira*, demonstrating that nitrification-related proteins of CMX *Nitrospira* populations were formed *in situ*. We found different genomic repertoires between CMX clade A and clade B affiliated *Nitrospira* MAGs containing genes for sulfur and alternative energy metabolism such as oxidation of formate and hydrogen, nitrogen transporter types and cellular regulation of reactive oxygen species. CMX clade B MAGs exclusively harbored genes involved in conversion of bicarbonate to carbon dioxide, oxidation of thiosulfate via the *sox* pathway, dissimilatory sulfate reduction, as well as genes encoding two superoxide dismutases (SOD1/ SOD2) and a formate transporter (*focA*). Transporters for cellular import of urea as well as for NO_2_ ^-^/NO_3_ ^-^ were present in representative genomes from both CMX clade A and clade B. We found that clade B MAGs encoded Amt-type NH_4_ ^+^ transporters, while clade A MAGs employed Rh-type NH_4_ ^+^ transporters. MAGs from both clades harbored genes for ATP citrate lyase (*acl*), which were transcriptionally active, indicating an active role in carbon fixation.

Compared to the CMX and AOA affiliated MAGs, transcription of *amo* genes was more than 100 times higher for AOB affiliated MAG H51-bin202-1 (Figure S 4). The AmoA sequence of this MAG had 93% sequence identity to the AmoA of the dominant, *N. ureae* affiliated AOB- OTU1 obtained from amplicon sequencing in this study (Figure S 5), suggesting that both approaches probably identified the same population of AOB. This MAG employed Rh-type NH4^+^ transporters as did representatives of CMX clade A, while AOA affiliated MAGs had Amt- type NH4^+^ transporters as representatives of CMX clade B (Figure S 4).

### Contributions of ammonia-oxidizing groups to ammonia oxidation activity

Short-term incubations in the presence of the nitrite oxidation inhibitor chlorate did not affect the ammonia oxidation rates (74.5 ± 9.4 nmol N L^-1^ d^-1^) compared to the rates in controls without inhibitor (75.9 ± 3.7 nmol N L^-1^ d^-1^) (Figure 4A). Chlorate is supposed to inhibit nitrite oxidation of NOB, and both ammonia oxidation and nitrite oxidation by CMX, as nitrite oxidoreductase catalyzes the formation of toxic chlorite from chlorate [7]. In the presence of the ammonia oxidation inhibitor allylthiourea, ammonia oxidation rates were not significantly different from zero. Allylthiourea is supposed to block ammonia oxidation, as it inhibits the ammonia monooxygenases of AOB as well as of CMX but affects those of AOA much less at the 100 µM concentrations used in this study [37, 60].

**Figure 4:**
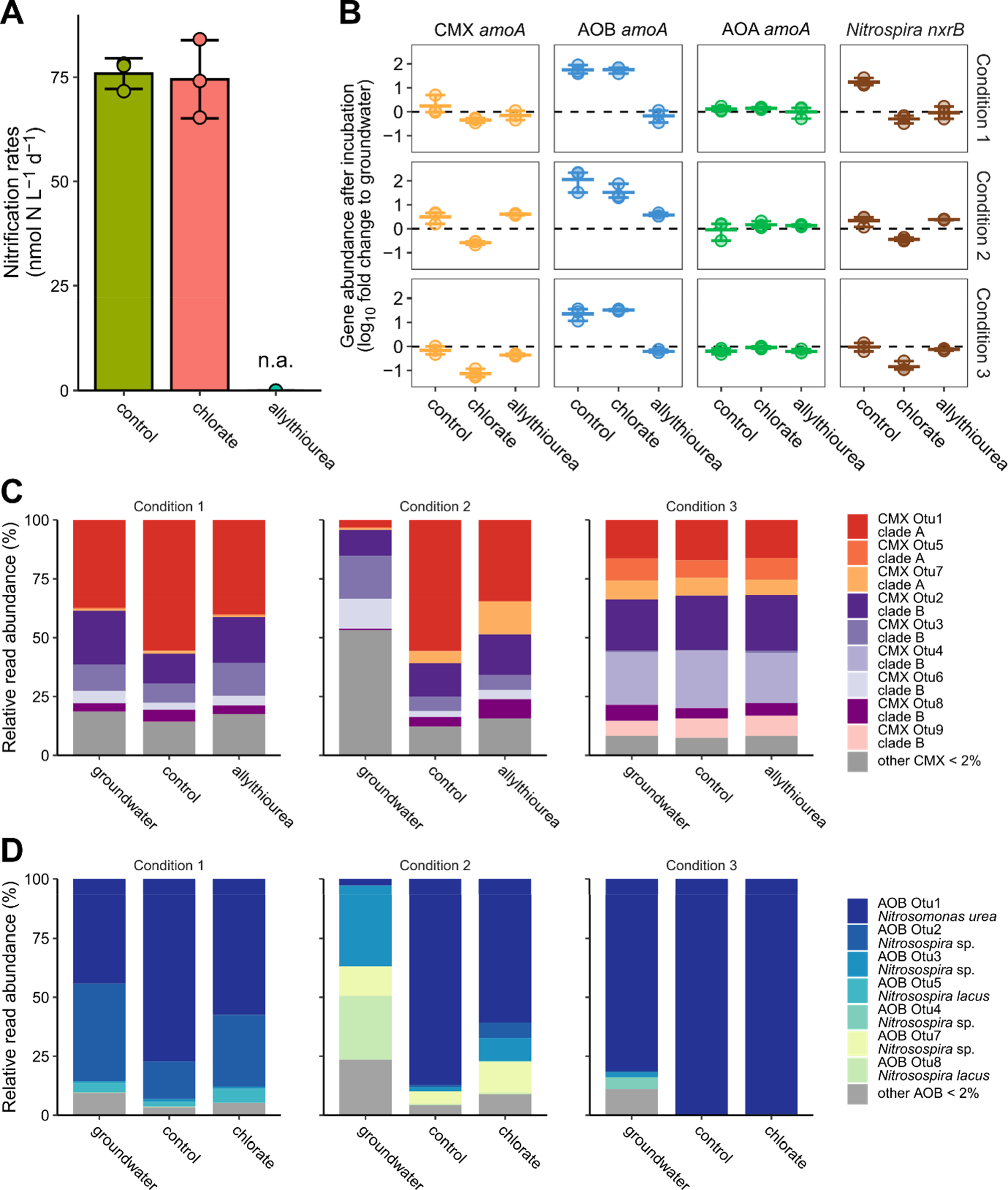
Groundwater incubations from wells H41 and H53 supplemented with different NH4^+^ concentrations and nitrification inhibitors across 5 days (short-term) and 25 to 35 days (long-term). **A)** Nitrification rates from short-term incubation of oxic groundwater supplemented with 50 µM NH4^+^ and selective inhibition with chlorate and allylthiourea (n.d. = not detected). Dots depict single rates of each replicate. Crossbar shows the average of three replicates and error bars show the variation between rates. **B)** Fold-change of *amoA* genes from ammonia oxidizers and *nxrB* genes from *Nitrospira*-like nitrite oxidizers after long-term incubation of groundwater. Single points represent one replicate and crossbar shows the average. Error bars display variance between conditions (n = 3). Condition 1: oxic groundwater supplemented with 10 µM NH4^+^. Condition 2: oxic groundwater supplemented with 50 µM NH4^+^. Condition 3: hypoxic groundwater without additional NH4^+^. **C)** Composition of CMX *Nitrospira* clade A and clade B affiliated OTUs of groundwater and after long-term incubation (n = 3). Bars show the mean relative *amoA* read abundance of OTUs representing > 2%. **D)** Composition of bacterial ammonia-oxidizing community and taxonomic affiliation of OTUs.

Long-term incubations (15-28 days) were performed to further investigate the potential growth and shifts in community composition of CMX, AOB, AOA and *Nitrospira*-affiliated NOB under the inhibitor treatments, using oxic groundwater from well H41 (condition 1: oxic/ 10 µM NH_4_ ^+^; condition 2: oxic/ 50 µM NH_4_ ^+^) and hypoxic groundwater from H53 (condition 3: hypoxic/ without supplement). Ammonia oxidation was always completely inhibited in the presence of allylthiourea, regardless of groundwater source or initial NH_4_ ^+^ concentrations (Figure S 6). NH_4_ ^+^ was only oxidized in incubations without inhibitor and in the chlorate treatment, while nitrite oxidation was only measured in incubations without inhibitor. Condition 3 without inhibitor showed complete consumption of NH_4_ ^+^ after 13 days, while it took 19 and 25 days under condition 1 and 2, respectively.

In incubations without inhibitor, CMX increased about two-fold (6.5 x10^5^ and 2.5 x 10^6^ *amoA* L^-1^ after 25 and 20 days of incubation) compared to initial groundwater (2.8 x 10^5^ and 1.44 x 10^6^ *amoA* L^-1^) under condition 1 and 2, respectively, while we did not detect growth of CMX under condition 3 (Figure 4B). CMX *amoA* gene abundance also increased in the allylthiourea treatment under condition 1 but decreased in the chlorate treatments. *Nitrospira*-affiliated NOB targeted using *nxrB* genes had similar growth patterns across condition and treatment types as CMX, and CMX accounted for an estimated 14 to 63 % of the *Nitrospira*-affiliated NOB population. AOB *amoA* abundance increased by one to two orders of magnitude in all untreated and chlorate treatments of the different conditions, while they decreased in the presence of allylthiourea. AOA *amoA* gene abundance showed only minor variations during all incubation settings. CMX OTU1 affiliated to CMX clade A (Figure 4C) and AOB-OTU1 related to *Nitrosomonas ureae* (Figure 4D) increased in abundance after incubations without inhibitor. These two OTUs were already dominant in the ammonia oxidizer communities of the original groundwater.

## Discussion

We present evidence that despite numerical predominance of CMX bacteria, they contribute only a small fraction to ammonia oxidation in the studied oligotrophic groundwater. Instead, ammonia oxidation in this system appeared to be largely driven by AOB. Ranging from 0.54 up to 136 nmol N L^-1^ d^-1^, the measured nitrification rates (ammonia oxidation + nitrite oxidation) matched those from oligotrophic lakes and open marine environments including oxygen minimum zones [61–64], but were lower than in rivers, and estuaries [65, 66] (Table S1). The observed nitrification rates did not reflect the spatial heterogeneity of groundwater NO_3_ ^-^ concentrations, suggesting that surface-derived inputs mask the effects of *in situ* NO3 formation on groundwater NO_3_ ^-^ loads in these carbonate-rock aquifers.

The oligotrophic conditions in the groundwater appeared to select for ammonia oxidizers with the highest reported NH3 affinities of their respective group. As the three groups differ in their NH3 affinity, with 0.049 to 0.083 µM for *Ca.* Nitrospira inopinata [15] (corresponding to 11 to 19 µM NH_4_ ^+^ at pH 7.3), 0.3 to 4 µM for the *Nitrosomonas oligotropha* lineage [15, 20] (corresponding to 55 to 829 µM NH_4_ ^+^ at pH 7.3), and < 0.01 µM for AOA closely related to *Nitrosoarchaeum koreensis* [15] (corresponding to < 2.3 µM NH_4_ ^+^ at pH 7.3), fluctuating resource availability at overall limiting conditions may have enabled their coexistence. Previous metagenomics analysis work at this study site identified CMX as key nitrifiers and demonstrated that measured groundwater carbon fixation rates could fully be accounted for by carbon fixation associated to nitrification, assuming nitrification and carbon fixation kinetics of CMX *Nitrospira* [21]. Similarly, in our study, CMX accounted for up to 80% of the ammonia oxidizer community at the groundwater well showing the highest nitrification activity, supporting our original assumption that CMX were the most competitive ammonia-oxidizing group. Our inhibitor-based approach strongly suggested that ammonia oxidation appeared to be largely driven by AOB, even though CMX outnumbered AOB by a factor of 10 in the samples used for rate measurements.

Treatment with 1 mM chlorate in nitrification assays was expected to inhibit the nitrite oxidation activity of canonical nitrite oxidizers as well as both steps of nitrification in CMX, since reduction of chlorate to toxic chlorite from the activity of nitrite oxidoreductase would eventually also inhibit ammonia oxidation in the same cell [7]. Ammonia oxidation was not affected in the chlorate treatment, but was completely inhibited in the treatment with allylthiourea, which specifically targets the ammonia monooxygenase and acts as a reversible inhibitor preventing the oxidation of NH_4_ ^+^ [60]. At a concentration of below 160 µM, it selectively inhibits the activity of AOB and CMX but not of AOA [37]. Consequently, this inhibitor approach strongly suggested that AOB were primarily responsible for the highest observed ammonia oxidation activity at well H41. A strong bias of our results by potential residual activity of CMX in the presence of chlorate appears unlikely. Although *Nitrospira* is capable of detoxifying hazardous chlorite by chlorite dismutases [67], the chlorate concentrations used here obviously resulted in growth inhibition and cell death, which is in line with previous observations in wetland sediment mesocosms [28]. Similarly, it appears unlikely that our results were caused by excessive growth of AOB, since nitrification rates were linear across these five-days incubations. A negligible contribution of AOA to ammonia oxidation was indicated by no detectable activity in the presence of allylthiourea.

We cannot completely rule out that activity by AOB was enhanced in the chlorate treatment due to absence of competition with CMX. This is one of the pitfalls associated with inhibition experiments and could have led to an overestimation of the contribution of AOB in our case. In fact, increased activity and growth of AOB in soil mesocosms where AOA were inhibited was recently demonstrated by Zhao *et al*. [68]. Our transcriptional and metaproteomics-based findings indicate a very active role of AOB in ammonia oxidation, supporting a substantial contribution of this group to ammonia oxidation rates. Ammonium concentrations at well H41, which had the highest nitrification activity, were around 10 µM NH_4_ ^+^ *in situ* and 50 µM were used for rate measurements, corresponding to 0.04 or 0.22 µM free NH3, respectively. The NH3 affinities and per cell ammonia oxidation rates differ between AOB and CMX [15]. It remains unclear to what extend competition for NH ^+^ would restrict their ammonia oxidation rates under the given ammonia-limited conditions. These conditions presumably affect AOB more than CMX, and if this would result in increased activity of AOB when CMX were suppressed due to inhibition. According to Vilardi *et al*., CMX and AOB can fill independent niches in the nitrifying community, which would reduce potential ammonia oxidation competition between both groups [69]. More detailed knowledge of the exact ammonia oxidation kinetics of the groundwater AOB and CMX representatives is needed to further disentangle the mechanisms of their coexistence.

Despite the presumably small contribution of CMX to ammonia oxidation rates, we obtained strong evidence of their active role in overall nitrification. CMX expressed genes and formed peptides involved in both ammonia and nitrite oxidation *in situ*, which was confirmed by growth of CMX in the presence of NH_4_ ^+^ in long-term incubations. However, it remains unclear if the major energy source to sustain cell metabolism originated from complete ammonia oxidation or rather from nitrite oxidation. In fact, a recent subsurface study detected transcriptional activity in association with nitrite oxidation but not ammonia oxidation for a CMX-related MAG [70], while CMX *Nitrospira* in water from rapid sand filters did not prefer to oxidize external nitrite alone [7]. As we found NO_2_ ^-^ transporters in representative groundwater MAGs from both CMX clades, the cells may also be capable of taking up the NO_2_ ^-^ formed by AOB, supporting their primary role as nitrite oxidizers along with *Nitrosomonas ureae* and *Nitrosospira* sp. affiliated AOB as key ammonia oxidizers.

AOB with faster ammonia oxidation rates than CMX [3, 15] did not outcompete CMX for NH_4_ ^+^ but simply processed a larger fraction of the available NH_4_ ^+^, thereby driving the ammonia oxidation rates and providing the resulting NO_2_ ^-^ to both CMX and canonical NOB. Overall, the genus *Nitrospira* clearly dominated the NOB community, as NOB affiliated with *Nitrospinaceae*, *Cand*. Nitrotoga, and *Nitrobacter* occurred only occasionally and were one or two orders of magnitude less abundant than *Nitrospira*. The question remains unclear, why CMX which accounted for more than 50% of the *Nitrospira* population were apparently favored as nitrite oxidizers over canonical nitrite-oxidizing *Nitrospira* in the groundwater. Physiological studies of *Ca.* Nitrospira inopinata demonstrated a much lower affinity to external NO_2_ ^-^ compared to canonical *Nitrospira* [15], a trait which is, however, not shared among all CMX *Nitrospira* [71]. A recent microcosm-based study of drinking water biofilter media suggested that CMX outcompete canonical NOB in natural nitrifier communities from nitrogen-limited environments over time [69]. However, co-occurrence of CMX and canonical *Nitrospira* appears to be common across a broad range of terrestrial and aquatic habitats [27], and their respective roles in nitrification might be highly variable.

In contrast to our findings, CMX *Nitrospira* were identified as the driver of the two steps of nitrification in groundwater-fed rapid gravity sand filters (RGSF) [7, 72]. Moreover, they made a similar contribution as AOB to ammonia oxidation in wetland soils [28]. At low NH_4_ ^+^ concentrations of 71 µM and 1.1 µmol g^-1^, CMX also dominated the ammonia oxidizer communities in the RGSF [7] and in coastal wetland sediments [28], respectively. However, our results demonstrate that high abundance of CMX in an environment does not necessarily imply that they also control the complete nitrification process. The competitive advantage of CMX has been associated with a biofilm-associated lifestyle such as in RGSF. However, division of labor in nitrification may remain the preferred strategy in the planktonic communities of groundwater environments at low NH_4_ ^+^ concentrations.

Distribution patterns of ammonia oxidizers across the groundwater wells suggested different niche preferences for AOB and CMX clade A compared to AOA and some clade B affiliated CMX. Both AOA and CMX clade B dominated in groundwater closer to the major recharge area with lower NH_4_ ^+^ availability. AOA can be the most abundant ammonia oxidizers in soils [73] and several studies reported CMX clade B in soil habitats [9, 74]. Consequently, transfer from soils to shallow groundwater could result in the observed high fractions of AOA and CMX clade B in the ammonia oxidizer communities of the respective wells [47]. In turn, deeper and more distant groundwater favored both AOB and clade A affiliated CMX under slightly increased availability of NH4^+^ and lower DO concentrations.

The observed ecophysiological niche differentiation of CMX clade A and clade B across different groundwater conditions was not reflected by differences in central nitrogen metabolism of representative MAGs [21, 30] and metaproteomic data from the groundwater. Genes encoding enzymes of ammonia and nitrite oxidation were highly transcribed and respective peptides represented both CMX clades. However, differences in the encoded NH_4_ ^+^ transporters could be linked to different ecological preferences. Rh-type NH_4_ ^+^ transporters in CMX clade A convey a better adaptation to higher NH_4_ ^+^ concentrations than Amt-type transporters. As Rh-type transporters have higher uptake capacity but lower NH3 affinity than Amt-type transporters [25] and are also encoded in AOB [75], CMX clade A might have a competitive advantage against clade B in the groundwater when more NH_4_ ^+^ is available. Along with NH_4_ ^+^ uptake efficiency, also oxygen tolerance appeared to drive the distribution patterns of the two clades, as CMX clade B but not clade A encode two types of superoxide dismutases (SOD1, SOD2) [25, 26]. Finally, given the broader metabolic versatility of clade B including the potential for formate and thiosulfate utilization, CMX clade B representatives might be less strongly tied to energy generation by nitrification compared to the clade A representatives.

In conclusion, our findings point to a strong contribution of AOB to ammonia oxidation activity in oligotrophic groundwater, despite numerical predominance of CMX. Given the limitations of differential inhibitor assays, the exact contribution of CMX to ammonia oxidation *in situ* remains unclear. Conditions of low NH_4_ ^+^ availability and varying oxygen concentrations appeared to generally support coexistence of CMX, AOB and AOA with representatives closely related to those with the highest NH_4_ ^+^ affinities in their respective groups. Our results suggest that high substrate affinity, high growth yields, and a potentially major role in nitrite oxidation enable CMX *Nitrospira* to maintain consistently high populations in the groundwater. We further propose niche differentiation of CMX clade A and B with a more important role in nitrification for clade A and a higher metabolic versatility, including the use of alternative electron donors, for clade B.

## Data availability

Sequence data generated in this study was submitted to the European Nucleotide Sequence Archive with project number PRJEB57321 for *amoA* genes under accessions ERS13665998 to ERS13665900, and with project number PRJEB63087 for bacterial 16S rRNA genes under accessions ERS15585018 to ERS15584991.

The mass spectrometry proteomics data have been deposited to the ProteomeXchange Consortium via the PRIDE [76] partner repository with the dataset identifier PXD039573.

Datasets containing the calculation of the groundwater nitrification rates, gene abundances, hydrochemical measurements and *amoA* OTU sequences will be released upon publication of the manuscript.

### Acknowledgements

This study was part of the Collaborative Research Centre 1076 AquaDiva (CRC AquaDiva) of the Friedrich Schiller University Jena, funded by the Deutsche Forschungsgemeinschaft under project number 218627073. Climate chambers to conduct experiments under controlled temperature conditions and the infrastructure for high-throughput sequencing were financially supported by the Thüringer Ministerium für Wirtschaft, Wissenschaft und Digitale Gesellschaft (TMWWDG; project B 715-09075 and project 2016 FGI 0024 “BIODIV”). Martin Taubert gratefully acknowledges funding by the Deutsche Forschungsgemeinschaft (DFG, German Research Foundation) under Germanýs Excellence Strategy – EXC 2051 – Project-ID 390713860.

We thank Heiko Minkmar, Falko Gutmann, Jens Wurlitzer, Stefan Krümmling, Swantantar Kumar, Patricia Geesink, Lena Carstens and Linda Gorniak for support of field and laboratory work and Gundula Rudolph for ion chromatography measurements. We acknowledge Kai Uwe Totsche and Robert Lehmann for field site management and provision of physicochemical data from groundwater.

## Author contributions

MH and MK designed the study. MK conducted all molecular work, incubation experiments and data analysis. BT and LB performed IRMS measurements and analysis. NC and WO performed metagenomic data analysis. MT, NJ and MvB conducted metaproteomic data analysis. MK and MH wrote the manuscript with contributions of all authors.

