## Supplemental Table 1 for "Differential contribution of nitrifying prokaryotes to groundwater nitrification"

#### Supplemental Material

Markus Krüger[1], Narendrakumar Chaudhari[1,2], Bo Thamdrup[3], Will Overholt[1], Laura Bristow[3], Martin Taubert[1], Kirsten Küsel[1,2], Nico Jehmlich[4], Martin von Bergen[2,4,5], Martina Herrmann[1,2*]

[1]Aquatic Geomicrobiology, Institute of Biodiversity, Friedrich Schiller University, Jena, Germany. [2]German Center for Integrative Biodiversity Research (iDiv) Halle-Jena-Leipzig, Leipzig, Germany. [3]Nordcee, Department of Biology, University of Southern Denmark, Odense, Denmark. [4]Department of Molecular Systems Biology, Helmholtz Centre for Environmental Research – UFZ, Leipzig, Germany. [5]Faculty of Biosciences, Pharmacy and Psychology, Institute of Biochemistry, University of Leipzig, Germany.

#### Supplemental Methods

##### $^{15}N$ rate incubations and IRMS analysis

Groundwater was sampled between 2018 and 2020 covering nine wells for rate measurements. Nitrification rate measurements of were conducted at a final concentration of 50 µM as $(^{15}NH_4)_2SO_4$. Samples were incubated at 15°C in the dark for five days without shaking. Subsamples of 10 ml were taken and replaced by equal amount of $N_2$ at the start of the experiment and after 12, 24, 48, 70 and 120 hours, followed by filtration through 0.2 µm filters and storage at -20°C until isotopic ratio mass spectrometry (IRMS) analyses. Nitrification rates were calculated based on the formation of $^{15}NO_2^-$ + $^{15}NO_3^-$ during incubation with $^{15}NH_4^+$. $^{15}NO_2^-$ and $^{15}NO_3^-$ were transformed to $N_2$ for analysis via cadmium reduction followed by a sulfamic acid addition [1, 2]. The $N_2$ produced ($^{15}N^{15}N$ and $^{14}N^{15}N$) was analyzed on a gas chromatography IRMS as previously described [3]. Single rates were determined from the slope of the linear regression of $^{15}N$ produced over time and corrected for the fraction of the $NH_4^+$ pool labelled in the initial substrate pool. Rates were assumed to be significantly different from zero, if the linear $^{15}N$ increase was significant ($p < 0.05$) (see Dataset S1). Abiotic $^{15}N$ transformation was excluded since filtered controls did not show any significant production of $^{15}N$ over the incubation period. The rate detection limit was 2.1 nmol N $L^{-1}$ $d^{-1}$ which was estimated from the median of the standard error of the slope from significant rates multiplied by the $t$ value for $p = 0.05$ [4]. This value estimates the magnitude of rates that may potentially escape detection. As the significance of rates is tested individually for each incubation, detectable rates may in some cases be below this detection limit.

##### Determination of inorganic nitrogen from mesocoms containing nitrification inhibitors

Subsamples for colorimetry of inorganic nitrogen compounds were taken at the onset and every two to three days until the end of incubation to monitor changes of the nitrogen chemistry. $NO_3^-$ from control and allylthiourea treated samples was measured using the sodium salicylate method [5]. As chlorate impaired the later $NO_3^-$ detection method [6], $NO_3^-$

from chlorate treated samples was determined via ion chromatography [6]. $NO_2^-$ and $NH_4^+$ were measured as indicated before.

**cDNA-synthesis and quantitative PCR reactions**

After extraction of total RNA from groundwater filters, 4 µl DNase treated RNA was used for reverse transcription in a total 15 µl reaction volume and 1 µl Random Primers following the manufacturer's protocol. Additional reactions without reverse transcriptase for each sample and a reaction with RT-PCR grade water instead of RNA were used to check for potential contamination with residual DNA in subsequent PCR steps. PCR using bacterial 16S rRNA primer pair 341f/785r [7] was performed to verify the presence of cDNA in the RT reactions and the absence of residual genomic DNA in the negative control reactions. The integrity of PCR products was checked by agarose gel electrophoresis.

Reactions for quantitative PCR were performed in 25 µl total volume with 12.5 µl Brilliant II SYBR Green qPCR Master Mix (Agilent), 0.4 µM of each primer and 5 µl of prediluted DNA or cDNA. Cycler conditions for *amoA* genes of AOB and AOA, and *nxrB* genes are described in [8] and [9], respectively. CMX *amoA* amplification was conducted in 45 cycles of 30 sec at 95°C, 45 sec at 52°C and 1 min at 72°C. Serial dilutions of standards used for gene quantification originated from plasmid DNA containing the respective gene as insert. Standard curves were linear from $5 \times 10^8$ to 50 copies per reaction with $R^2 > 0.99$ and efficiencies ranging from 80 to 95%.

Transcript/ gene ratios of CMX, AOB and AOA were calculated by dividing the *amoA* gene abundances $L^{-1}$ by the *amoA* transcripts $L^{-1}$. To estimate the proportion of CMX *Nitrospira* to canonical *Nitrospira*, the percentages of *amoA* and *nxrB* abundance were calculated from the sum of both gene abundances at each groundwater well. *Nitrospira nxrB* abundance was divided by 1.5 to account for a higher copy number estimated from *nxrB* copy numbers from Hainich *Nitrospira* MAGs.

**Amplicon sequencing of *amoA* genes and sequence analysis**

PCR for amplicon library construction was performed in 50 µl reactions including 2 µl of DNA, 0.4 µM of each primer, 1 µg/ µl BSA and 2X HotStartTaq Master Mix (Qiagen). PCR conditions for AOB *amoA* were 35 cycles for 45 sec at 94°C, 30 sec at 57°C and 45 sec + 1 per cycle at 72°C and final 10 min at 72°C. AOA *amoA* was amplified using 35 cycles for 45 sec at 94°C, 60 sec at 53°C and 60 sec at 72°C, and final 10 min at 72°C. For amplification of CMX *amoA* the following protocol was used: 40 cycles for 30 sec at 95°C, 30 sec at 53°C and 1 min at 72°C ending with 10 min at 72°C [10]. The size and integrity of all PCR products was validated by agarose gel electrophoresis. Paired-end 2 x 300 bp reads were sequenced on an Illumina MiSeq instrument using v3 chemistry [11].

Primer sequences of raw *amoA* reads were trimmed with Trim Galore (https://github.com/FelixKrueger/TrimGalore). Sequences were quality filtered with DADA2 [12] using filterAndTrim default settings and truncLen of 200, 229 and 260 bp for CMX, AOB and AOA, respectively. Subsequent steps were performed in Mothur [13] including merging of paired reads and dereplication, followed by chimera search using the uchime algorithm implemented in Mothur v.1.46.1 [14]. Reads were translated to amino acid sequences with transeq incorporated in Emboss [15]. Sequences containing stop codons were excluded from further analysis. The remaining reads were assigned to OTUs using vsearch at 95% for CMX, 95% for AOB [16] and 96% for AOA [17] sequence identity. OTUs with < 10 reads were removed before classification. The taxonomical assignment of *amoA* sequences was done with BLASTx [18] using a custom *amoA* database on amino acid level and identification of closest relatives with best hits. AmoA sequences of OTUs accounting for more than 2% of all reads were aligned using Clustal Omega [19]. Phylogenetic tree construction of deduced AmoA amino acid sequences was performed using maximum likelihood with the JTT+CAT evolutionary model and 1000 bootstraps in FastTree [20] and trees were visualized in iTOL [21].

**Inferring metabolic capacities of nitrifying prokaryotes from metagenome assembled genomes**

In addition to sorted CMX *Nitrospira* MAGs, other nitrifier MAGs from Overholt et al. [22] which employed either *amoA* for AOB and AOA and *nxrA* or *nxrB* for NOB affiliated genomes were included in the analysis. Genes encoding for key pathways involved in nitrogen and alternative energy metabolism and uptake, as well as genes for carbon fixation and cell defense mechanisms against reactive oxygen species were examined.

**Protein extraction and mass spectrometric analysis**

For metaproteomic analysis, tryptic peptides were dissolved in 0.1% formic acid (v:v) and subjected to LC-MS/MS analysis on a Q Exactive HF instrument (Thermo Fisher Scientific, Waltham, MA, USA) in LC chip coupling mode, equipped with a TriVersa NanoMate source (Advion Ltd., Ithaca, NY, USA). Raw mass spectral data was analysed with the Sequest HT search algorithm in Proteome Discoverer (v1.4.1.14, Thermo Fisher Scientific, Waltham, MA, USA). The following parameters were used: enzyme specificity was set to trypsin with two missed cleavages allowed, carbamidomethylation (cysteine) was set as static modification and oxidation (methionine) as dynamic modification, and peptide ion and MS/MS tolerances were set to 10 ppm and 0.05 Da, respectively. Peptides were considered identified upon scoring a q-value < 1% based on a decoy database and obtaining a peptide rank of 1.

**Statistical analysis**

Statistical analysis was performed in R 4.12 (R Core Team, 2020) using packages FSA [24], ComplexHeatmap [25] and vegan [26]. Shapiro-Wilk test was applied to test for normal distribution of data before conducting either parametric or non-parametric tests. Significant differences of groundwater hydrochemical properties and gene abundances between groundwater wells were determined using Kruskal-Wallis rank sum test and Dunn's multiple comparison post-hoc test with package FSA [24]. Correlations between hydrochemical properties, total *amoA* abundances and absolute *amoA* read abundances of OTUs were calculated using Spearman's rank correlation and heatmap construction with

ComplexHeatmap package [25]. Ordination analyses were performed with vegan [26]. For redundancy analysis, the absolute *amoA* read abundances of OTUs were standardized using Hellinger method [27] and hydrochemical parameters were log-transformed before performing rda function from vegan. Nonmetric multidimensional scaling using Bray-Curtis distance matrix was performed to show dissimilarity of ammonia oxidizing communities between groundwater from different wells. Anosim function from vegan was used to verify the significance of observed differentiation.

**Figure S1**

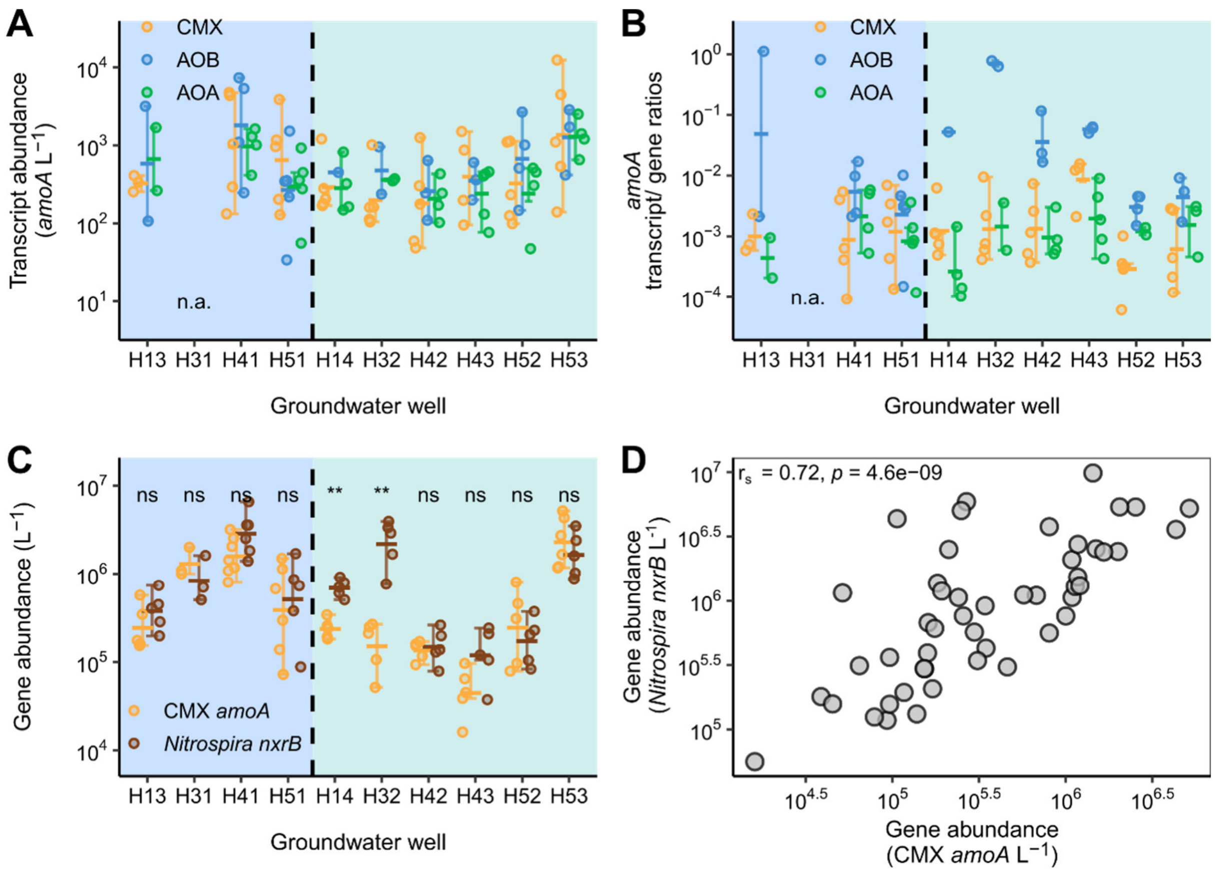


Figure S 1 Display of **A)** *amoA* transcripts $L^{-1}$, **B)** *amoA* transcript/ gene ratios and **C)** CMX *amoA* and *Nitrospira*-like *nxrB* genes $L^{-1}$ across groundwater sites (n ≥ 3). Dots represent single sample measurements (n.a. = no RNA sample due to low sample volume), the crossbar shows the mean of all samples and the error bars display the variation from the median. Outliers are shown beyond the error bars. Lowercase letter code indicates significant differences based on Dunn's multiple comparison test (* = $p < 0.05$, ** = $p < 0.01$, ns = not significant). **D)** Correlation between CMX *amoA* genes and *Nitrospira*-like NOB *nxrB* genes across all groundwater wells based on Spearman's rank correlation.

**Figure S2**

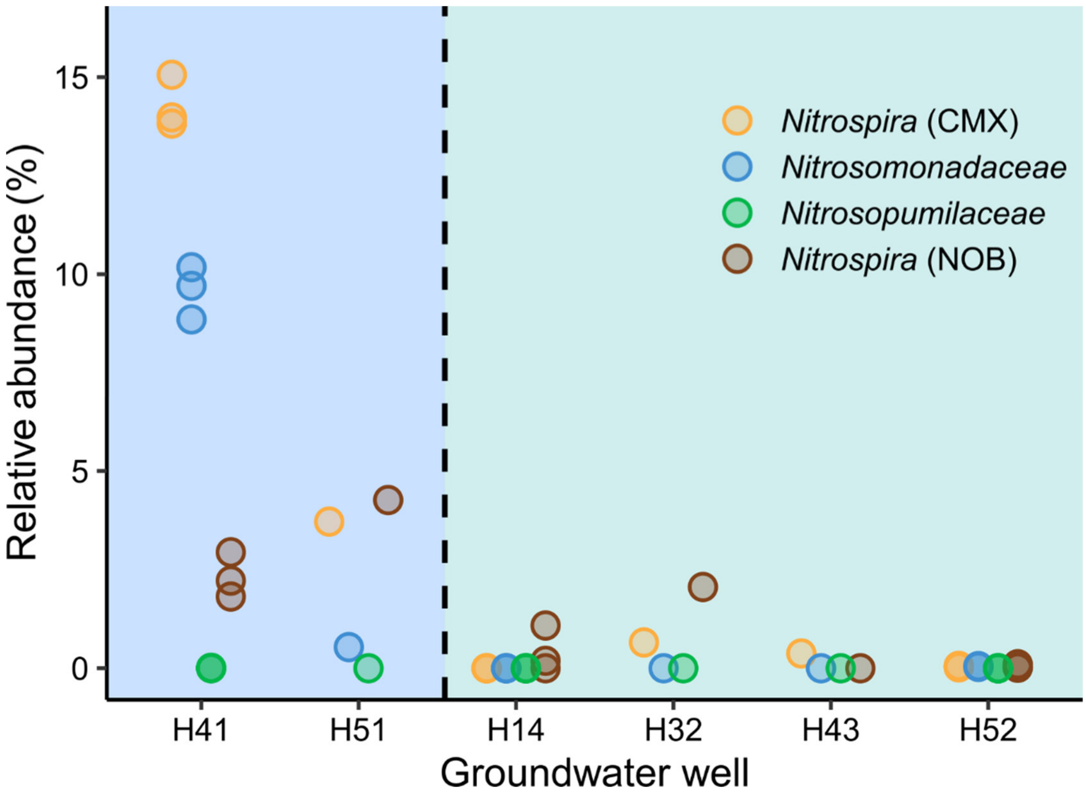


Figure S 2: Relative abundance of peptides in relation to whole peptide abundance per sample. Single dots represent MAGs affiliated with *Nitrosomonadaceae*, *Nitrosopumilaceae*, CMX *Nitrospira* and canonical *Nitrospira* derived from a metaproteomic data set.

142 **Figure S3**

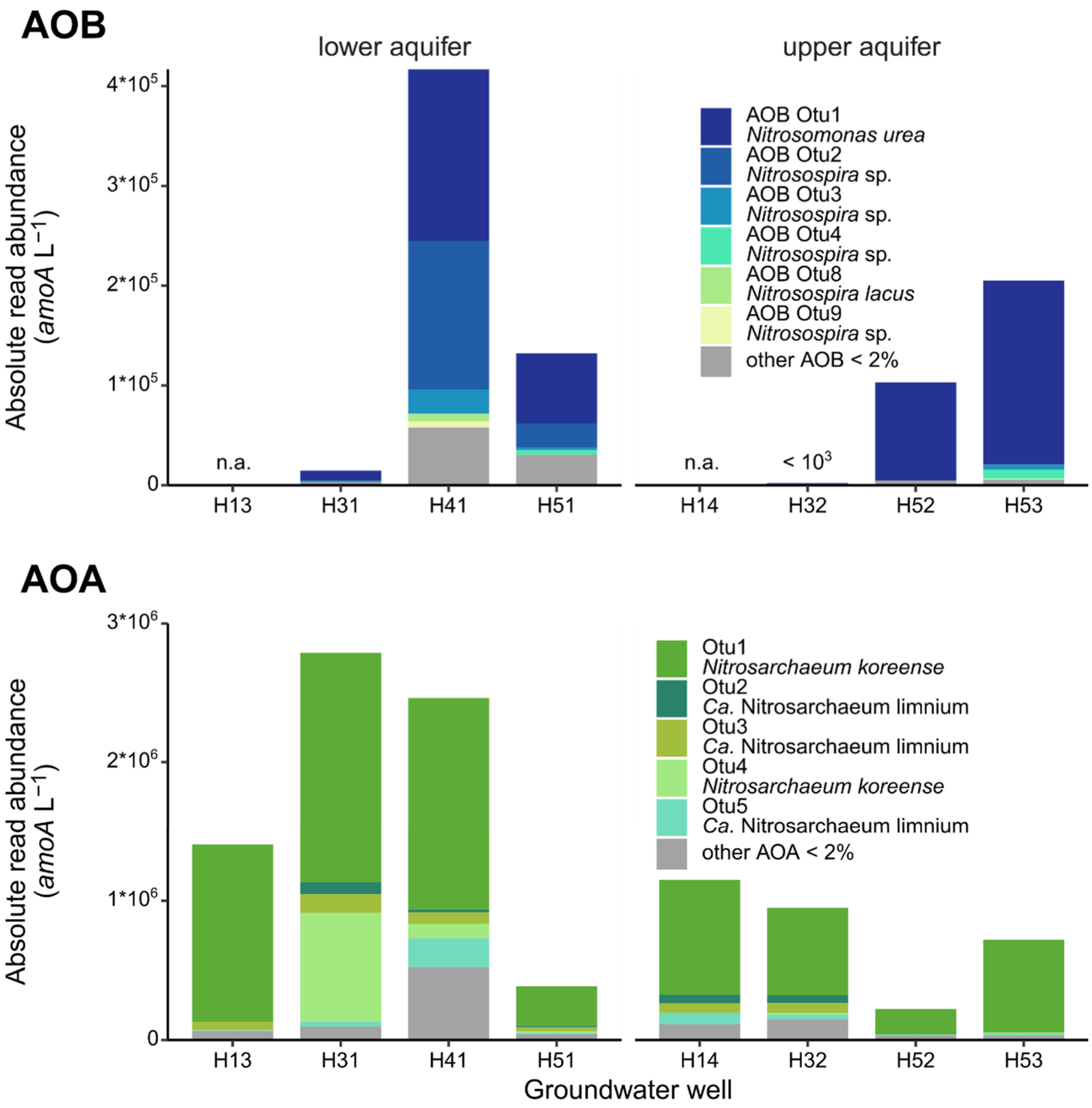


143

144 Figure S 3: Composition of the groundwater AOB and AOA community, and taxonomic
145 affiliation of OTUs across eight groundwater wells (n ≥ 3 per well). Each bar displays the mean
146 absolute *amoA* read abundance of OTUs representing > 2% of each ammonia oxidizer
147 community (n.a. = not sequenced due to low abundance).

**Figure S4**

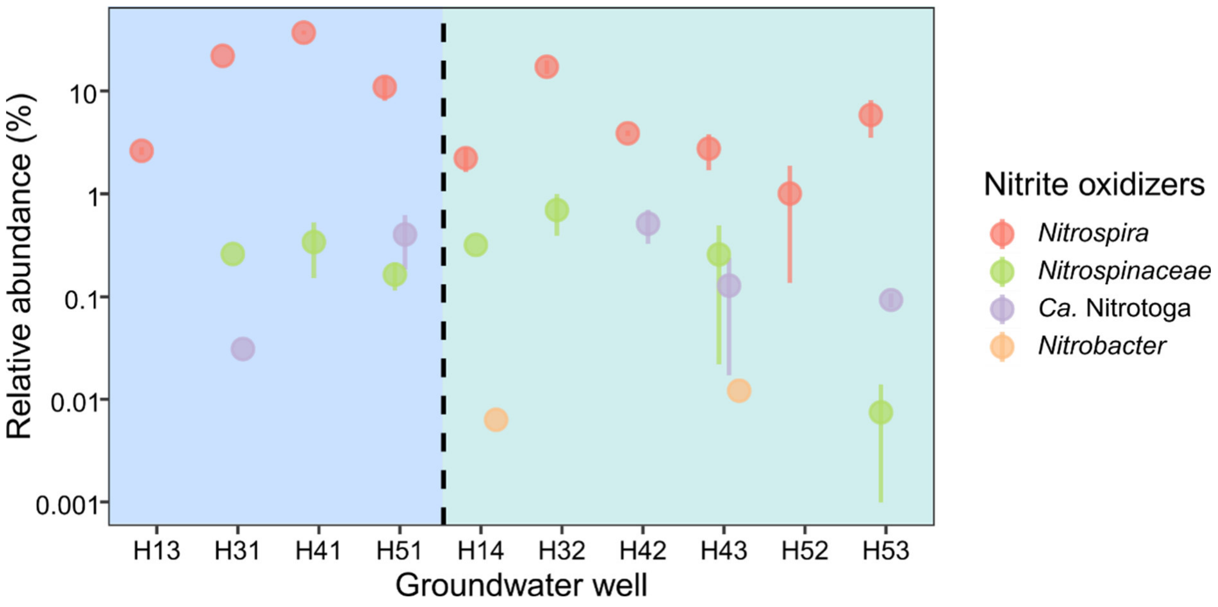


Figure S 4: Relative abundance of 16S rRNA genes affiliated with different bacterial nitrite oxidizers across the groundwater wells.

### Figure S5

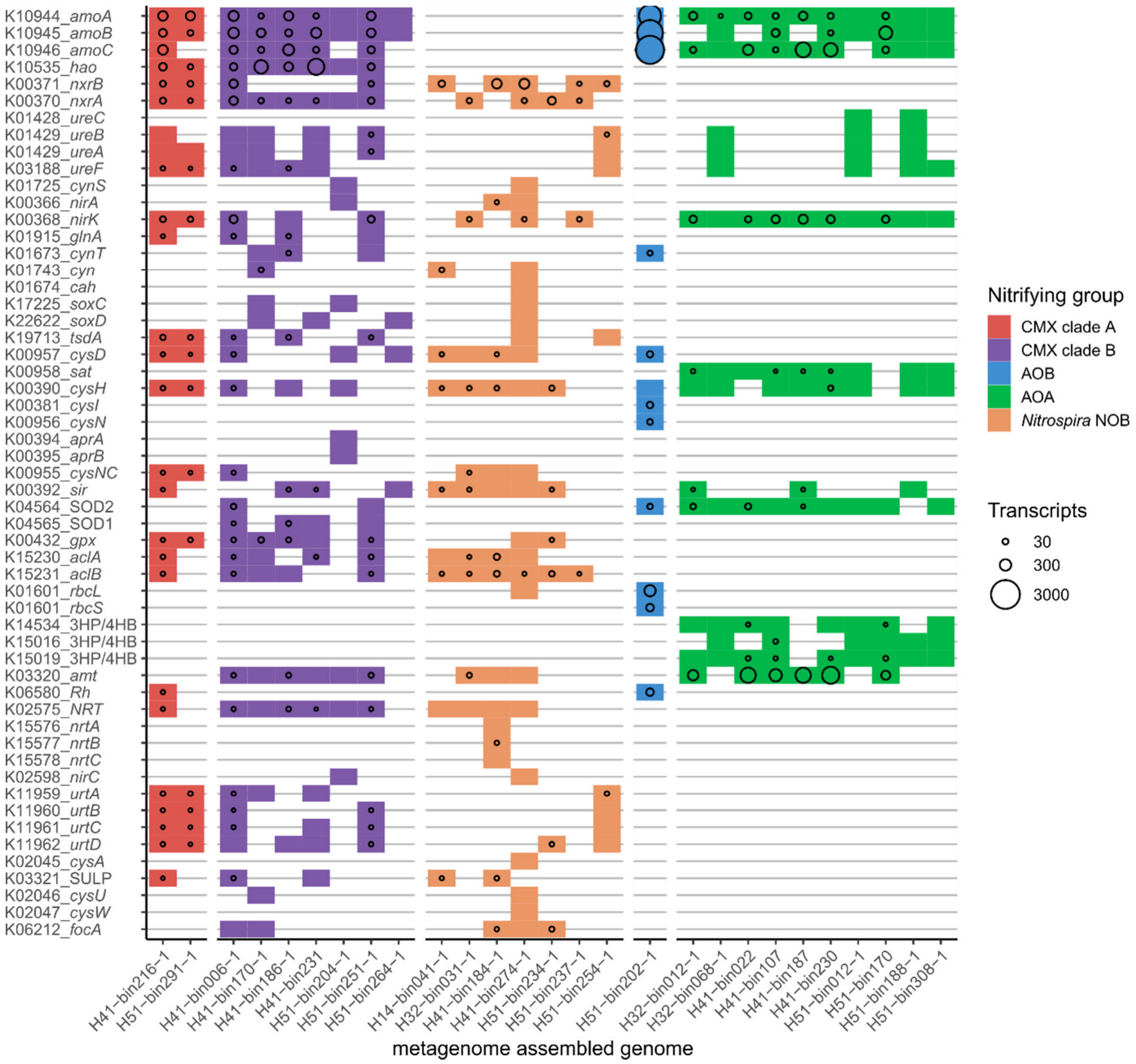


Figure S 5: Genomic potential in nitrogen and sulfur energy metabolism as well as protection against radical oxygen species and transporters of Hainich groundwater MAGs affiliated with CMX *Nitrospira* clade A and clade B, *Nitrospira*-like NOB, AOB and AOA. The size of circles displays the maximum transcript coverage for each gene derived from metatranscriptomic data.

**Figure S6**

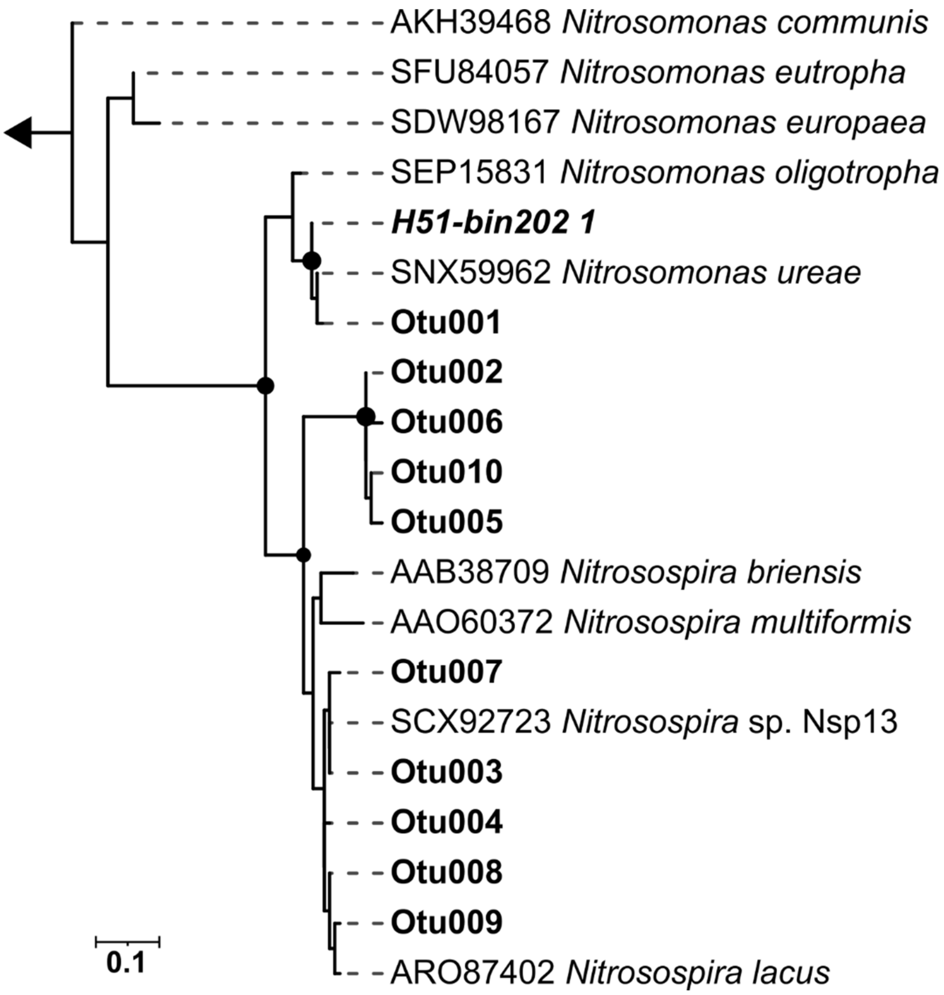


Figure S 6: Maximum likelihood tree of deduced AmoA amino acid sequences from ammonia oxidizing bacteria consisting of ten OTU amplicon sequences (bold), one *Nitrosomonas* MAG sequence (bolditalic) and nine reference sequences. The tree was constructed using JTT substitution model with gamma distribution and 1000 bootstrap iterations. Bootstrap support values greater than 75% are shown as black dots. Two *amoA* sequences from *Nitrosococcus* were used as an outgroup indicated by the arrow.

### Figure S7

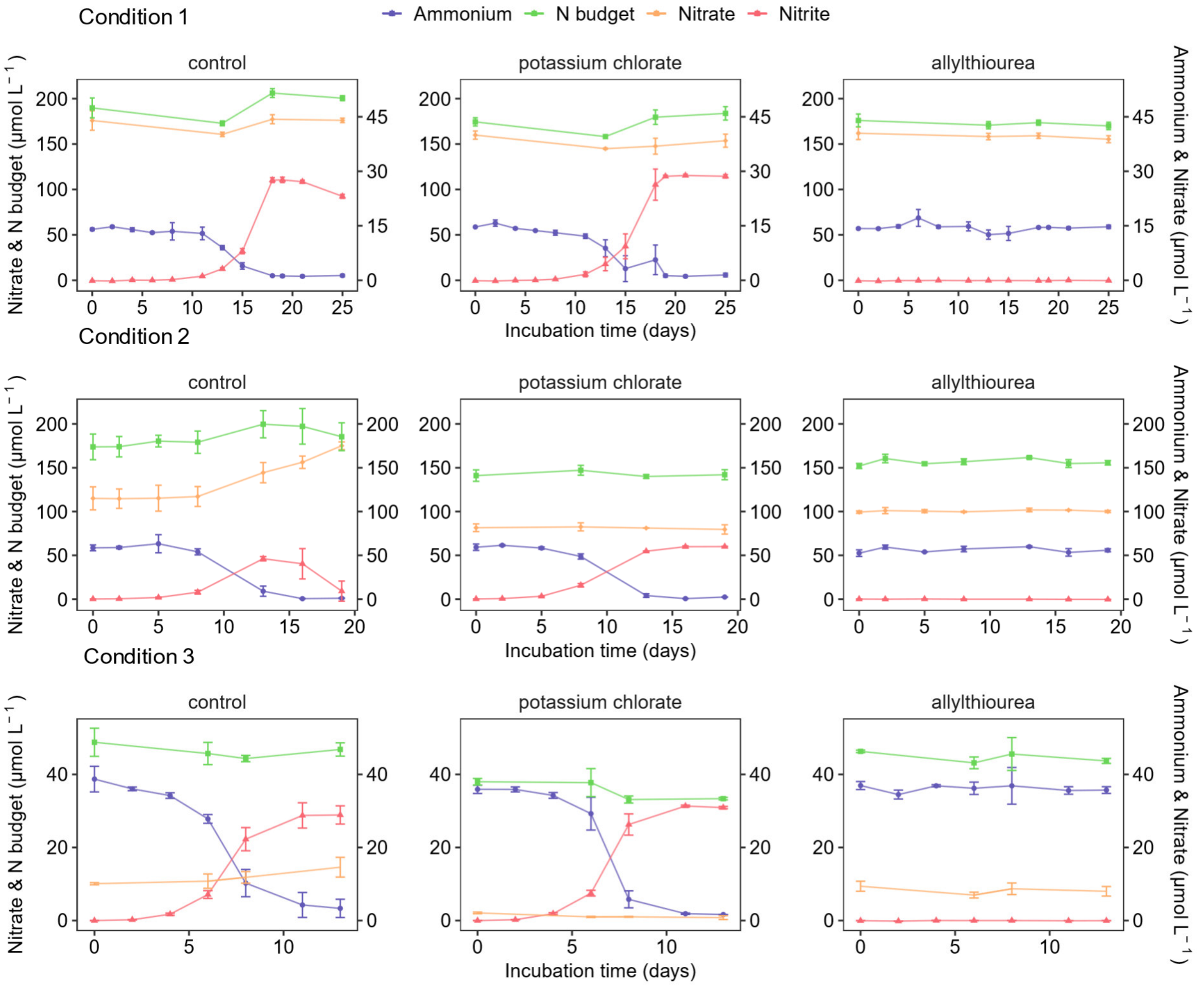


Figure S 7: Changes in the concentrations of ammonium, nitrite and nitrate in groundwater incubations in the presence of nitrification inhibitors. Incubations were conducted with groundwater from oxic well H41 (condition 1: 10 µM $NH_4^+$, condition 2: 50 µM $NH_4^+$) and hypoxic well H53 (condition 3: no supplement). Two different inhibitors were chosen to selectively target groups of ammonia oxidizers (potassium chlorate inhibits CMX, allylthiourea inhibits AOB and CMX) and controls without inhibitor. Ammonium, nitrite and nitrate were monitored during incubation of groundwater. Each dot displays the mean of three replicates and standard deviation is visible where larger than symbol. N budget represents the accumulated concentrations of ammonium, nitrite and nitrate in each incubation. Black arrows indicate addition of 15 µM $NH_4^+$ after 15 days of incubation.

#### Table S1

Table S 1: Overview of ammonium oxidation rates across different environments based on $^{15}N$ labelling approaches.

| **Environment** | **Depth m** | **$NH_4^+$ Ox nmol N $L^{-1}$ $d^{-1}$** | **Reference** |
|---|---|---|---|
| carbonate-rock aquifers | 5 - 88 | 0.54 – 136.3 | this study |
| oligotrophic freshwater lake | 2 - 80 | 1.8 - 50.5 | Small *et. al.* 2013 [28] |
| freshwater marsh | < 2 | 9,600 - 16,800 | Gribsholt *et. al.* 2005 [29] |
| river estuary | < 1 | 1.44 - 3,984 | Miranda *et. al.* 2008 [30] |
| marine oxygen minimum zone | 40 - 50 | 2.4 - 40 | Bristow *et. al.* 2016 [4] |
| Arabian sea | 50 - 136 | 1.8 - 22.5 | Newell *et. al.* 2011 [31] |
| San Pedro Ocean Time-series SPOT | 45 | 13.8 – 40.2 | Beman *et. al.* 2011 [32] |
| Bermuda Atlantic Time-series BATS | 150 | 8.13 – 16.7 | Beman *et. al.* 2011 [32] |
| HOT oligotrophic North Pacific | 175 | 1.61 – 8.14 | Beman *et. al.* 2011 [32] |
| Sargasso Sea oligotrophic | 240 | 1.01 – 1.58 | Beman *et. al.* 2011 [32] |

#### Supplemental Datasets

Dataset S1: Calculation of groundwater nitrification rates and inhibitor approaches using linear regression and determination of limit of detection.

Dataset S2: Abundances of *amoA*, *nxrB* genes and transcripts across the groundwater wells and incubation experiments determined by qPCR.

Dataset S3: Taxonomic classification of ammonia oxidizer OTUs generated by *amoA* targeted amplicon sequencing of groundwater samples.

Dataset S4: Specifications of Hainich nitrifier MAGs and reference genomes.

16. Purkhold U, Wagner M, Timmermann G, Pommerening-Röser A, Koops HP. 16S rRNA

and *amoA*-based phylogeny of 12 novel betaproteobacterial ammonia-oxidizing isolates: Extension of the dataset and proposal of a new lineage within the nitrosomonads. *Int J Syst Evol Microbiol* 2003; **53**: 1485–1494.

24. Ogle D, Wheeler P, Dinno A. FSA: Fisheries Stock Analysis. R package version 0.8.31. 2020.
